# Defining the architecture of cerebrospinal fluid cellular communities in neuroinflammatory diseases

**DOI:** 10.1101/2021.11.01.466797

**Authors:** Tina Roostaei, Claudiu Diaconu, Hanane Touil, Claire Harbison, Ya Zhang, Samantha Epstein, John Tuddenham, Kiran Thakur, Julien Bryois, Heinz Wiendl, Gerd Meyer Zu Hörste, Dheeraj Malhotra, Claire Riley, Vilas Menon, Philip L. De Jager

## Abstract

Cerebrospinal fluid (CSF) biomarkers are important for multiple sclerosis (MS) diagnosis. Moreover, absent of autopsy or biopsy tissue, CSF is the most relevant source for studying the immune cells involved in MS pathophysiology. Single-cell RNA sequencing (scRNA-seq) provides new opportunities to advance our understanding of disease-associated changes in CSF immune cells. Here, using scRNA-seq data generated from 58 CSF and 10 PBMC samples, we provide an updated atlas of the immune cells present in human CSF in MS and other neuroinflammatory conditions, including novel lymphoid and myeloid cell clusters. Our atlas can thus serve as a reference for future studies of immune cells in neuroinflammation. Our further characterization of CSF myeloid cells suggests that most CSF microglia-like cells resemble two of the previously-described brain microglia signatures. Additionally, our data from a sex-mismatched bone marrow transplant recipient suggest that CSF microglia-like cells are of peripheral origin. Our comparisons between MS and other neuroinflammatory disorders show a highly-specific increase in plasma cells, along with reductions in the proportion of microglia-like cells in MS CSF. Furthermore, our analyses on MS patients receiving anti-CD20 therapy ocrelizumab suggest that the treatment effects are not limited to B cell depletion, and ocrelizumab appears to reverse some MS-associated T and myeloid changes in CSF. Finally, we utilized our atlas to prioritize (1) CSF cell types expressing genes associated with MS susceptibility, and (2) ligand-receptor gene pairs that are differentially expressed in MS CSF, providing targets for further mechanistic and causal investigations in pathophysiology and treatment of MS.

## Introduction

Multiple sclerosis (MS) remains a diagnosis of exclusion today. One integrates the clinical history with magnetic resonance imaging and targeted biofluid (blood and cerebrospinal fluid, CSF) analysis to evaluate the likelihood that this inflammatory and neurodegenerative disease is present. Given the utility and less-invasive nature of imaging and peripheral blood measurements, many practitioners treat the examination of CSF as optional. However, one cannot deny the value of additional information gathered from CSF analysis, which is often reserved for ambiguous cases. CSF provides information on the compartmentalized central nervous system pathology that is not measurable using other data sources (e.g., the presence of oligoclonal bands). Moreover, its assessment using new technologies offers great potential for advancing our understanding of the immune portion of the disease pathophysiology and identifying new targets for treatment in studies of living humans.

Single-cell RNA sequencing (scRNA-seq) studies have in the recent years started to characterize the CSF immune cells in a number of disease conditions^1–4^. In contrast to flow cytometry studies that rely on information collected on a pre-defined list of proteins, scRNA-seq studies work with information collected on the whole transcriptome of single cells. As a result, scRNA-seq studies have not only been successful in providing complementary information on the diversity of CSF lymphoid and myeloid cells, they have also been fundamental in the discovery of previously undescribed cell types, e.g., a group of microglia-resembling myeloid cells that are present in the CSF regardless of diagnosis, and are absent from the peripheral blood.

Here, taking advantage of a substantially larger sample size and a variety of diagnoses, we provide a fine-grained characterization of CSF immune cells commonly found in MS and neuroinflammatory conditions. Our CSF cell map contains detailed subclassification of the lymphoid and myeloid, including the microglia-like cells, and reliably covers smaller cell clusters with 1:1000 frequency in CSF. Our study design is further utilized to (1) provide insight into the ontogeny of CSF microglia-like cells, (2) prioritize immune cell types and genes relevant to early MS in comparison to neuroinflammation of other causes, and (3) assess the cellular and molecular effects of B cell depleting therapy on immune cells in MS CSF. Overall, our neuroinflammation CSF cell map can serve as a reference for future studies of CSF, and our cell composition and gene expression findings can be leveraged for further mechanistic investigations into MS onset and treatment.

## Results

### Description of participants

We recruited two sets of participants from the Columbia University MS Center and the Clinical Neuroimmunology Service at Columbia University Irving Medical Center: (1) individuals undergoing a diagnostic lumbar puncture (LP) as part of an initial clinical evaluation for suspected central nervous system (CNS) inflammation and (2) individuals with progressive multiple sclerosis (PMS) treated with ocrelizumab (OCR) who underwent LP solely for research purposes. Participants in the OCR group had all been on ocrelizumab treatment for >2 years and were sampled approximately 5 months after their last infusion; all were clinically and radiographically stable. At the conclusion of the diagnostic process for the first group (patients with suspected neuroinflammatory disease), a final diagnosis was established for each patient. All patients diagnosed with remitting-relapsing MS (RRMS) were grouped together. These patients were all untreated within at least 1 month of the LP and had a gadolinium-enhancing lesion on MRI within 3 weeks of the LP. The other patients had a variety of diagnoses and were grouped into the “other neuroinflammatory disorders” (ONID) category (**Supplementary Table 1**). All, except 2 of the ONID patients also had at least one gadolinium-enhancing CNS lesion within 3 weeks of the LP, except for a participant with isolated optic neuritis and a participant with leukoencephalopathy. All were untreated within 1 month of LP, except for a participant with lymphoma who presented with acute encephalitis and had been treated with methylprednisolone at the time of LP. One of the patients in the ONID group underwent a second research LP after 2 months, and therefore had two separate samples. Demographic details are presented in **Supplementary Table 1** (n=31 samples from 30 participants).

CSF processing was performed as outlined in the **Supplementary Methods**; in short, CSF was centrifuged at 300g for 10 minutes at 4°C to collect the cell pellet which was resuspended in 70μL of CSF prior to loading on the Chromium platform from 10x Genomics. All samples were profiled immediately after collection, and they were all loaded on the Chromium platform within 4 hours of CSF extraction.

To enhance the power of our study in detection of less frequent cell types, determining the CSF-specificity of cell clusters, and assessing the MS-specificity of our findings, we repurposed previously reported^3,4^ CSF (n=22) and PBMC (n=10) data from untreated RRMS and idiopathic intracranial hypertension (IIH) patients collected at the time of diagnostic LP at the Department of Neurology at the University of Münster. We also included additional CSF (n=5) data from individuals undergoing diagnostic LP at the University of Basel who were later diagnosed with IIH or ONID. The final number of samples included in the study from the three centers was 68 (58 CSF samples from 57 individuals, in addition to 10 PBMC samples) (**Figure 1a**). All demographic and clinical details are presented in **Supplementary Table 1.**

**Figure 1.**
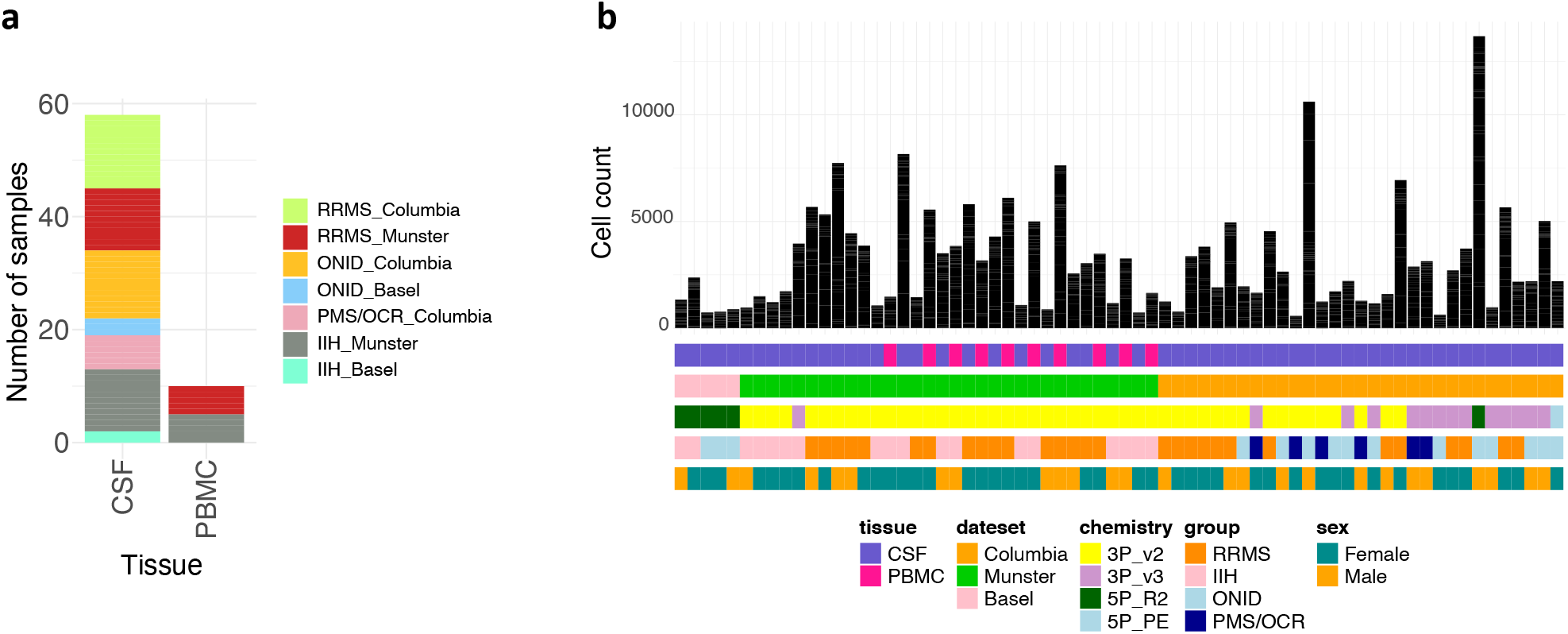

### Description of the single cell data and clustering structure

Data were generated from each sample using one of four different chemistries based on the Chromium platform. All samples were processed using a single analytical pipeline, and each sample was individually preprocessed before integration. Briefly, doublets were identified using Scrublet^5^, each sample was clustered using default Seurat^6^ parameters, and RBC clusters were identified. These steps were followed by the removal of doublets, RBC clusters, and additional low-quality cells (i.e., cells with >25% mitochondrial transcripts or <100 unique identified genes) (more details in the **Supplementary Methods**). Of note, in order to avoid biased exclusion of cells based on their characteristics, we only excluded cells that were not of interest, like RBCs, and extremely low-quality cells that did not have enough features for proper dimensionality reduction and clustering. Background removal was then performed using SoupX^7^. All samples had at least 500 cells after QC. The distribution of each sample’s final cell counts and characteristics can be found in **Figure 1b**. We also performed an initial annotation of cells on a cell-by-cell basis by mapping each sample’s filtered data to the Azimuth PBMC atlas^8^ using supervised PCA^8^, in addition to mapping to a number of other atlases that contain immune cell RNA expression data using SingleR^9^.

After the initial quality control and individual sample preprocessing, 216,723 transcriptomes (175,529 CSF cells and 41,194 PBMC) from 57 individuals were integrated into a common space using Harmony^10^. Given that each sample underwent sequencing in a separate run, cells from each sample were considered to be of a different batch. Integration was set to account for the effects of batch, chemistry, and collection site (**Supplementary Figure** ◻). The integrated data were then used for an initial clustering effort to identify the major cell types in our data. Manual annotation of the clusters using cluster marker genes combined with Azimuth annotations for cells in each cluster identified several T, Natural Killer (NK) and myeloid cell clusters, in addition to a large cluster of B cells, a plasma cell cluster, a cluster of proliferating cells, and a few other PBMC-dominant clusters such as hematopoietic stem cells and platelets (**Figure 2a**). Of note, we found two sample-specific T cell clusters (clusters 17 and 21) that were not grouped with the larger connected population of T and NK cells. These two clusters belonged to the CSF sample from the participant with ◻ lymphoma and expressed *CXCL13*. The pathology report for this CSF sample had noted the presence of atypical mature T cells. Hence, it is likely that the two clusters capture the mentioned atypical T cells which are not found in other samples.

**Figure 2.**
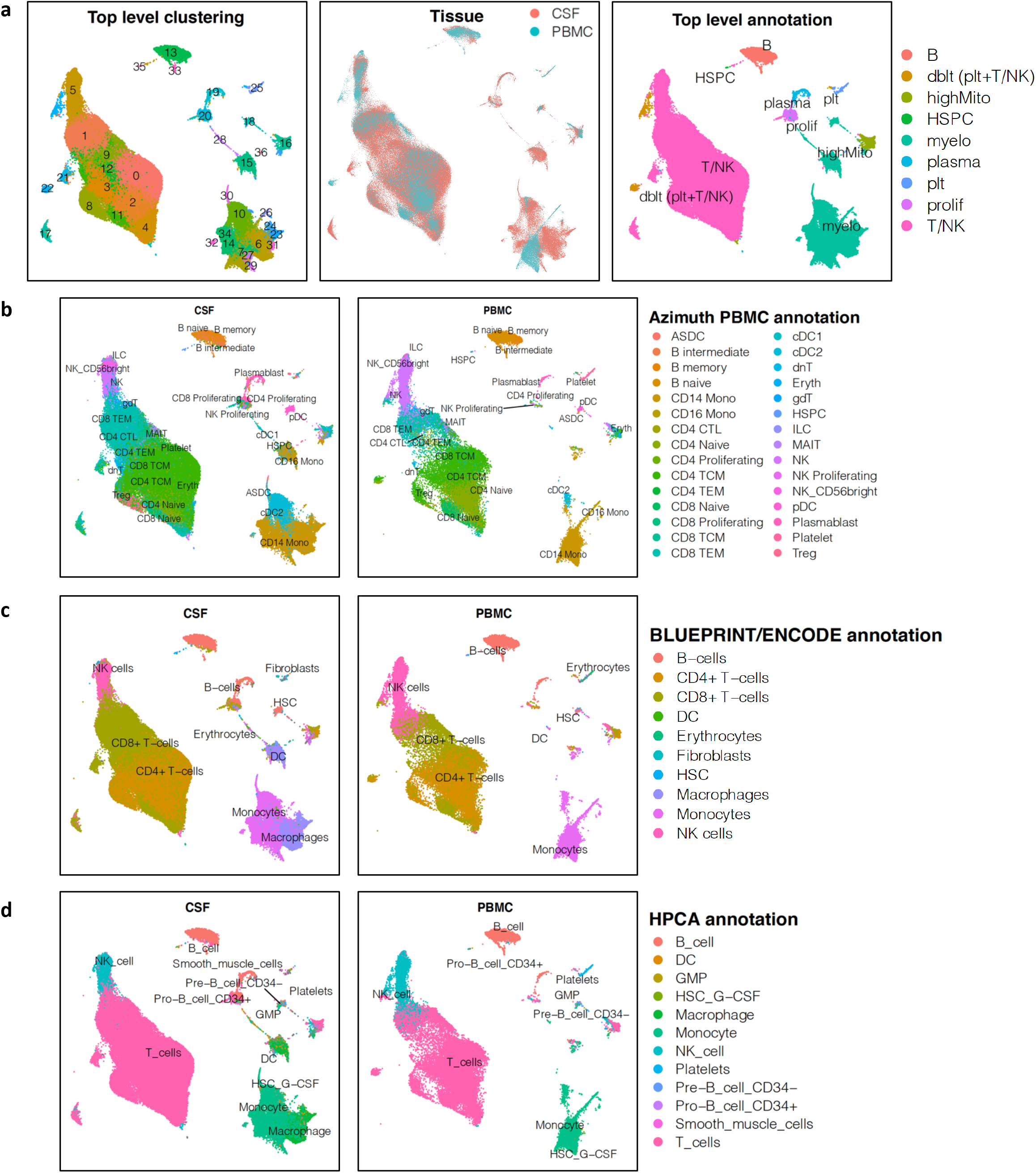

Among the myeloid cell populations we noted several clusters of monocytes, dendritic cells and microglia-like cells. The microglia-like clusters expressed previously-identified marker genes such as *C1QA/B/C*, *APOE*, and *APOC1*. These cells were annotated as monocytes when aligned to Azimuth PBMC data (**Figure 2b**). However, mapping them to transcriptome data from the Human Primary Cell Atlas^11^ (HPCA) and BLUEPRINT/ENCODE^12,13^ projects (which include data from additional immune and non-immune cell types that are not present in PBMC) identified CSF microglia-like cells as macrophages (**Figure 2c-d**). Mapping cells to these two atlases further suggested that no major granulocyte cluster was present in the data, as is expected since PBMC isolation removes the granulocytes. Having identified the major cell types in our samples, we then turned to the second stage of our clustering effort in which we separated the major cell types and reclustered each cell type to enhance the resolution of cell subtypes.

### Secondary clustering of T and NK cell populations

As seen in **Figure 3a**, our secondary clustering of 167,599 T and NK cells (140,679 cells from CSF and 26,920 cells from PBMC) resulted in 20 clusters. Similar to the initial clustering, we annotated the identified clusters using the cluster marker genes and the annotations from the Azimuth PBMC atlas. The two sample-specific and likely-atypical CXCL13^+^ T cell clusters from the participant with lymphoma still formed distinct clusters and were not studied further. In our higher resolution model, the large population of NK cells were more clearly separated from the T cells. However, similar to the results from previous scRNA-seq studies, the rest of the T cells were clustered close together, suggesting that they have less distinguishable features at the RNA level and might be better defined as a continuum. This observation is in line with previous studies where scRNA-seq data is shown to be suboptimal in identifying known subclasses of T cells that are classically defined by protein expression data^8^.

**Figure 3.**
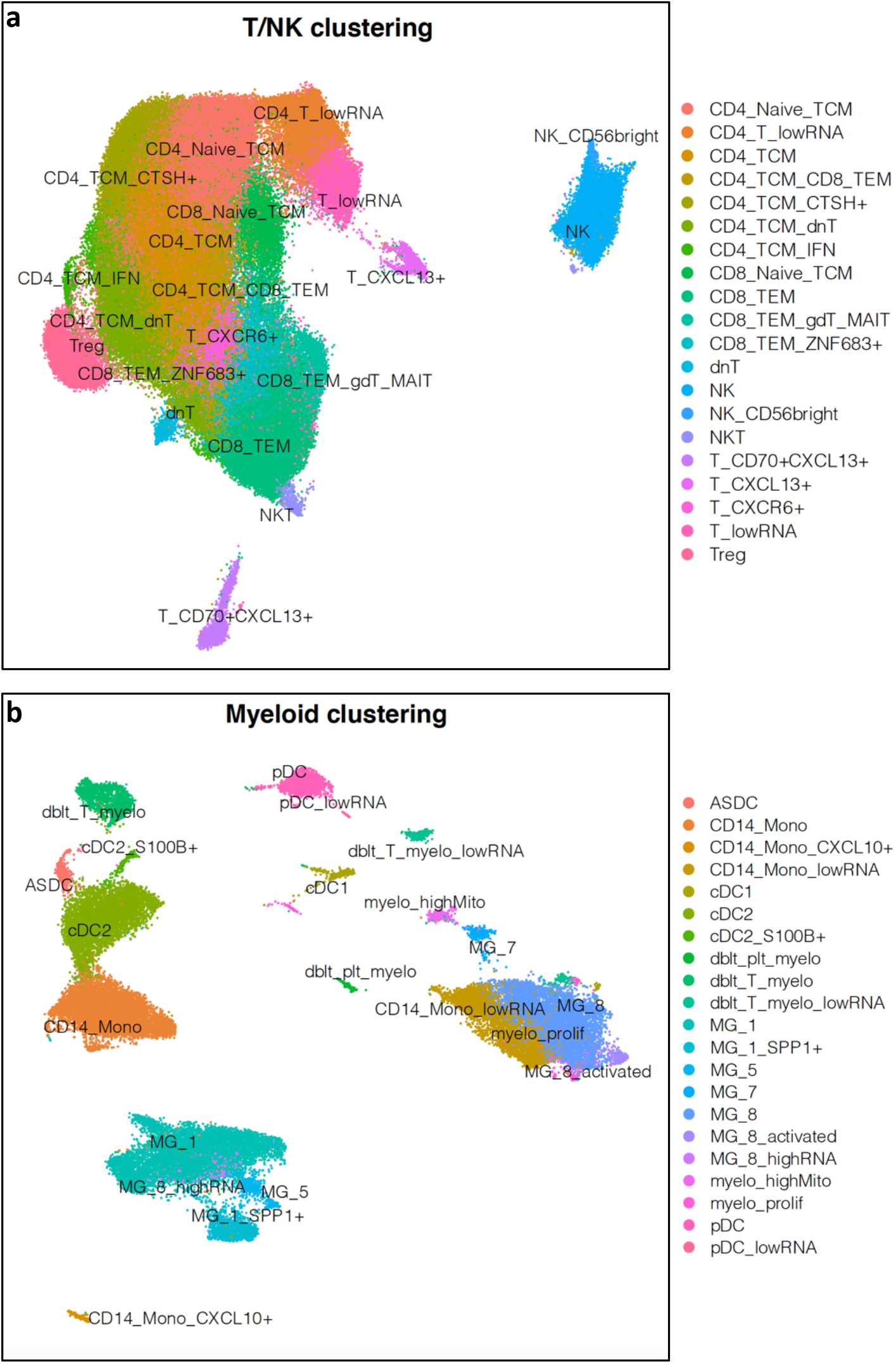

Among the large group of connected T cell clusters, we identified clusters that were dominated by one or few of the classic T cell subtypes, such as naïve or memory CD4^+^ or CD8^+^ T, regulatory T, double-negative T, gamma-delta T, mucosal associated invariant T (MAIT) or NKT cells. However, a subset of our clusters could more specifically be defined by the expression of genes not previously known as marker genes for functional T cell subtypes: *CTSH* was expressed in a cluster of CD4^+^ memory T cells; *ZNF683* expression distinguished a cluster of CD8^+^ memory T cells; a CD4^+^ memory T cell cluster expressed several interferon-induced genes including *ISG15*, *IFI44L*, *IFI6*, *IFIT3*, and *IFIT1*; and finally, we had a cluster of *CXCR6*^+^ T cells, of which >99% of its cells were from CSF samples. *CXCR6* is a marker of tissue-resident memory T cells (TRM), and the CSF-specificity of this cluster suggests that cells captured in this cluster might be long-lasting memory T cells that have spent a substantial time in the CNS parenchyma and are transitioning back into the peripheral circulation. The majority of cells in this cluster were annotated by Azimuth as CD4^+^ (62%), followed by a smaller proportion of CD8^+^ T cells (21%). To confirm our interpretation, we took a recently reported^14^ signature of TRM cells from non-CNS tissues and assessed for enrichment of the TRM RNA signature in all T cell clusters. Our CXCR6^+^ T cell subset is the one that is most enriched for the TRM signature (**Supplementary Figure** ◻). Since this CXCR6^+^ T cell cluster is specific to the CSF, it may therefore be of particular interest in translational studies as a useful surrogate for monitoring immune responses within the CNS parenchyma. Notably, as seen in the next sections, this T cell subtype is one of the T cell clusters that its frequency is altered in MS in comparison to other neuroinflammatory disorders.

In addition to the more classic T cell clusters and the clusters distinguished by their relatively novel markers, we found two other clusters than contained 10% of cells in the T cell component. These clusters had slightly less RNA content in comparison to the rest of the T cell clusters (**Supplementary Figure** ◻), and showed fewer identifying features. One of these clusters was dominated by CD4^+^ naïve and memory T cells, and the other was dominated by CD4^+^ and CD8^+^ memory T cells. Cells from these clusters were not sample- or tissue-specific, and although we cannot rule out the possibility of technical or artifactual effects, their large proportion might well suggest a biological underlying effect such as a not-yet-characterized functional or metabolic T cell state.

Additional mapping of our T and NK cells to bulk RNA-seq samples of sorted immune cell populations from the Monaco Immune Data^15^ and Database of Immune Cell Expression^16^ (DICE) (**Supplementary Figure** ◻) suggested that a majority (73%) of the cells in the CD4^+^CTSH^+^ TCM cluster express a Th1/Th17 or Th17 transcriptomic signature. This observation is in line with the literature on *CTSH* expression as a marker for both Th1/Th17 and Th17 cells^17^. Additionally, a substantial proportion of cells in the CD4^+^ TCM/dnT (20%) and CD4^+^ naïve/TCM (15%) clusters appear to express a signature close to follicular helper T cells.

### Secondary clustering of myeloid cell populations

As seen in **Figure 3b**, our secondary clustering of 36,463 myeloid cells (28,289 cells from CSF and 8,174 cells from PBMC) resulted in 21 clusters. Using marker genes and Azimuth, HPCA and BLUEPRINT/ENCODE atlases, we annotated several clusters of dendritic cells (DC), which include the classical (cDC1 and cDC2), plasmacytoid (pDC), and AXL^+^SIGLEC6^+^ (ASDC) subtypes coming from both CSF and PBMC samples. Interestingly, a novel cluster of CD1c^+^ dendritic cells emerged in the secondary clustering that expresses both *S100B* and *ACY3*. This small cluster contained 0.6% of myeloid cells and mainly consisted of cells from CSF samples (98%), raising the possibility that they may reflect a subset involved in activating the CXCR6^+^ TRM cells in our samples from individuals with neuroinflammation.

In order to better characterize the macrophage/microglia-like clusters, we took advantage of our recent scRNA-seq atlas of human microglia purified from cortical tissue samples collected at autopsy or surgery^18^. We combined data from the microglial atlas with data from another scRNA-seq atlas of sorted peripheral blood monocytes and dendritic cells^19^ in a SingleR analysis in order to map our myeloid cells to a relatively comprehensive set of characterized scRNA-seq reference myeloid data from both blood and brain. Our analysis suggested that the CSF microglia-like clusters resemble specific subtypes of human microglia. The majority of microglia-like cells are found in two large clusters: one mostly resembles MG-1 cells, which are believed to be homeostatic microglia; and the other expresses an MG-8 signature, which is found in microglia cells with lower RNA content and downregulated marker genes, potentially representing a different metabolic or functional state. In addition to these two larger clusters, 3 of the 5 remaining microglia-like clusters were also dominated by MG-1- and MG-8-like cells: a cluster of MG-1-like cells that also expressed *SPP1*, a cluster of MG-8-like cells that expressed *CD9* and *HSPB1*, likely representing an activated state, and a sample-specific cluster of MG-8-like cells with high RNA content from the sole participant with rheumatoid meningitis. The remaining 2 microglia-like clusters, on the other hand, were either dominated with cells resembling MG-5, which is a subtype of cytokine-producing microglia, or MG-7, which have heightened expression of genes involved in antigen-presentation and may represent a class of professional antigen presentation cells. These results expand the narrative of the presence of microglia-like cells in the CSF to include the fact that they exist in several distinct subtypes, reflecting different functional specifications. Of note, a proportion of PBMC monocytes grouped with microglia-like clusters in our secondary clustering. The exceptions were the MG-5 and MG-1-SPP1^+^ clusters that seemed to be CSF-specific.

Contrary to the dendritic and microglia-like cells that mostly got re-distributed over clusters of the same cell class in our second stage of clustering, monocytes got re-distributed over both monocyte-dominant and microglia-like dominant clusters in the secondary clustering. PBMC CD16^+^ monocytes, which formed a separate cluster in the initial clustering, were clustered with CSF MG-1 cells. PBMC CD14^+^ monocytes were found in 3 major clusters: 75% of them were equally distributed over the two final CD14^+^ monocyte clusters, one of which had lower RNA content and might represent a transitional state, while another 17% were clustered with CSF MG-8 cells. CSF clearly contained both types of CD14^+^ monocytes. Additionally, we observed a small cluster of CD14^+^ cells consisting of both CSF and PBMC cells, which expressed *BIRC3* and *CXCL10*, and might represent a transitional state with a signature close to mature dendritic cells^20^.

The projection of clusters from the secondary clustering of T, NK and myeloid cells onto the initial cell map for the CSF samples can be seen in **Figure 4**.

**Figure 4.**
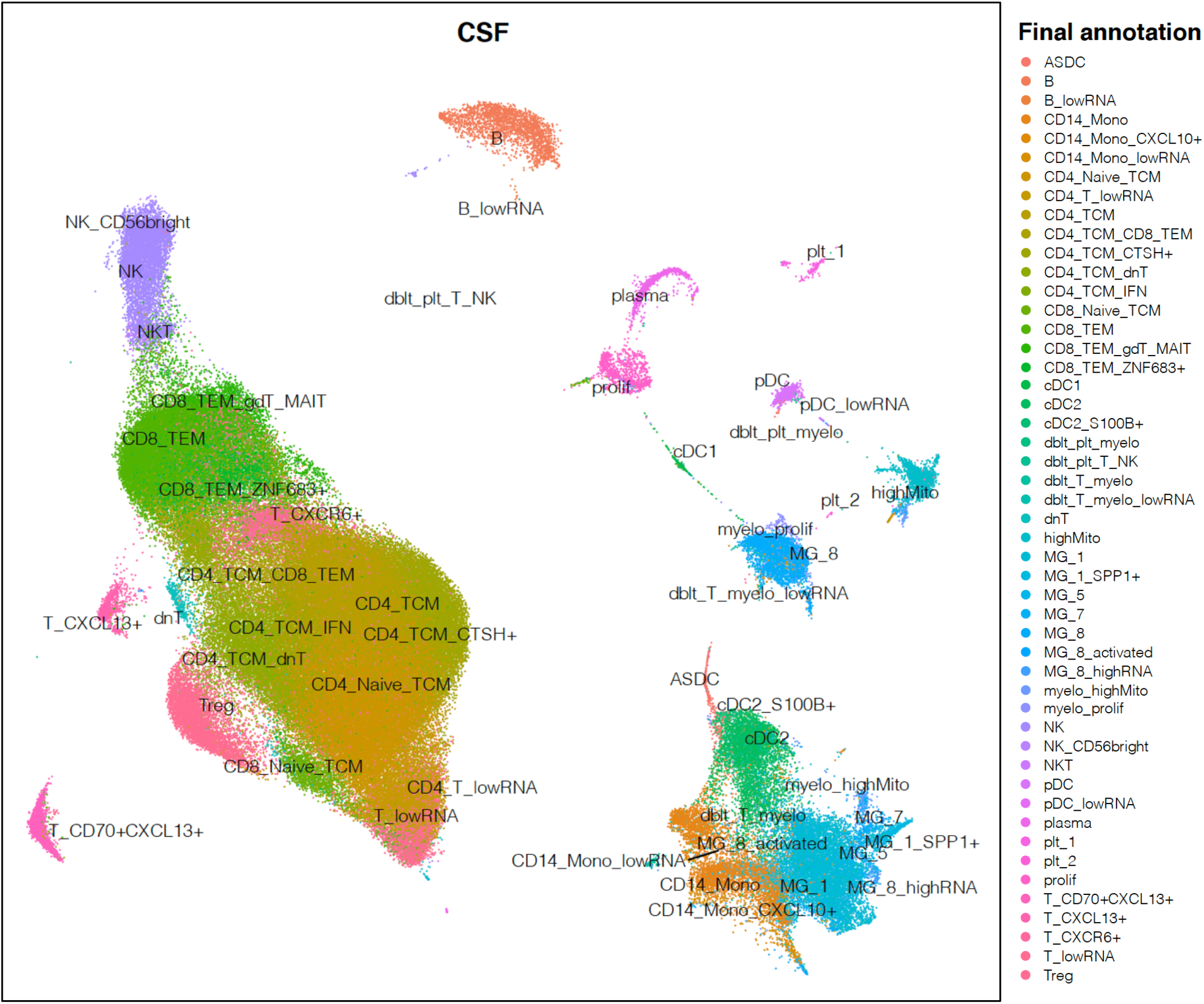

### Origin of CSF microglia-like cells

Animal studies have provided evidence that meningeal and perivascular macrophages are derived from the bone marrow, while microglia are derived from yolk sac progenitors^21^. These findings are corroborated by human brain studies in sex-mismatched bone marrow transplant recipients^22^. In order to address the question of the origin of the CSF microglia-like cells, we took advantage of scRNA-seq data from one of our participants from the ONID group: she is a female individual with chronic progressive multifocal leukoencephalopathy (PML) arising from infection of the brain with the John Cunningham (JC) virus in the setting of a bone marrow transplant performed 10 years earlier for ◻ lymphoma. The transplant occurred after a course of ◻ chemotherapy and was from a male donor. CSF samples were collected at two different time points, 2 months apart, and profiled with scRNA-seq as part of the presented experimental and analytic pipeline. The frequencies of the identified cell types were highly comparable between the two time points (Spearman’ρ =0.93), and the samples contained cells from all major CSF cell types, including microglia-like cells (**Supplementary Figure** ◻). RNA expression of sex chromosome genes suggested that all cell clusters from both samples had an attributed male origin (**Supplementary Figure** ◻). Moreover, deconvoluting genetic identity of the cells using Freemuxlet^23^ showed that all cells were from a single origin. These data suggest that CSF microglia-like cells are unlikely to be emigrating microglial cells; instead, they are likely to be derived from the bone marrow and to adopt microglial-like features while they are in the CNS. However, their exact cells of origin and their differentiation process remains unclear.

### Enrichment of CSF cell types for MS susceptibility genes

The functional consequences of most MS susceptibility variants (defined using our latest genome-wide map of MS susceptibility^24^ with its 234 variants) remain poorly understood. We therefore took the opportunity to see whether certain CSF cell types or subtypes might be enriched for MS susceptibility genes since these cells are particularly relevant to the disease. Our analysis using this list of variants and the MAGMA^25^ method suggests that the following CSF cell types as enriched for genes associated MS susceptibility loci: B and plasma cells, naïve and memory CD8^+^ T cells, NKT, NK, Treg, dnT, MG-1 and MG-5. Interestingly, no CD4^+^ T cell subset and most of the other microglia-like and dendritic cell and monocyte subtypes were not enriched in MS susceptibility genes using this method (**Supplementary Figure** ◻). We also used a different approach, using the list of 558 prioritized MS susceptibility genes that we shared in our report describing the 234 variants^24^, and we find the same cell subtypes to be enriched, along with a few additional myeloid and CD4^+^ T cell subtypes (**Supplementary Figure** ◻).

### Can CSF cell composition distinguish RRMS from other neuroinflammatory disorders?

Identifying immunopathological differences that can distinguish between MS and other neurological and neuroinflammatory conditions can not only help with diagnostic challenges but also advance our understanding of the mechanisms underlying the disease and potentially identify targets for intervention. In the next set of analyses, we leveraged our defined CSF cell population structure to identify cell types whose frequencies are altered in RRMS in relation to ONID (Columbia samples). We then performed a similar analysis comparing RRMS and IIH (Munster samples), and utilized these results to prioritize the more specific cell type changes in MS by looking for cell types with altered frequencies in both sets of comparisons.

Starting with major cell types, we observed an increase in the proportion of T and plasma cells, and a decrease in the proportion of myeloid cells in RRMS compared to ONID (all adjusted *p*<0.05, **Figure 5a**). Similar pattern was observed for these cell types in comparison with IIH (trend significance for T cells, *p*=0.06, and significant differences in myeloid and plasma cells, *p*<0.05, **Figure 5b**). The observed T-lymphocytic predominance in MS CSF is in line with previous cytometric studies of MS and ONID[ref]. We were also able to replicate our finding on increased lymphoid frequency in MS in an independent set of CSF samples assessed using clinical cytometry measures (24 MS and 23 ONID patients evaluated by the Clinical Neuroimmunology Service at Columbia University Irving Medical Center in 2019-2020, **Figure 5c, Supplementary Table** ◻). Despite significant differences, the proportions of T and myeloid cells are overlapping among different conditions, and neither of these cell population frequencies provide high diagnostic value for MS. The ratio of T:myeloid cells[ref] has the same challenge (**Figure 5**). On the other hand, the proportion of plasma cells shows more promise, as plasma cells appear to be almost absent from non-MS CSF (**Figure 5a**). Interestingly, our finding on this cell type aligns well with results from the few flow cytometry studies^26–28^ that have assessed plasma cells or plasmablasts in MS CSF compared to other neurological diseases. Consistent with a recent observation^26^, a cut-off of 0.25% for plasma cells showed 100% specificity and 70% sensitivity for distinguishing RRMS from ONID and IIH in our samples.

**Figure 5.**
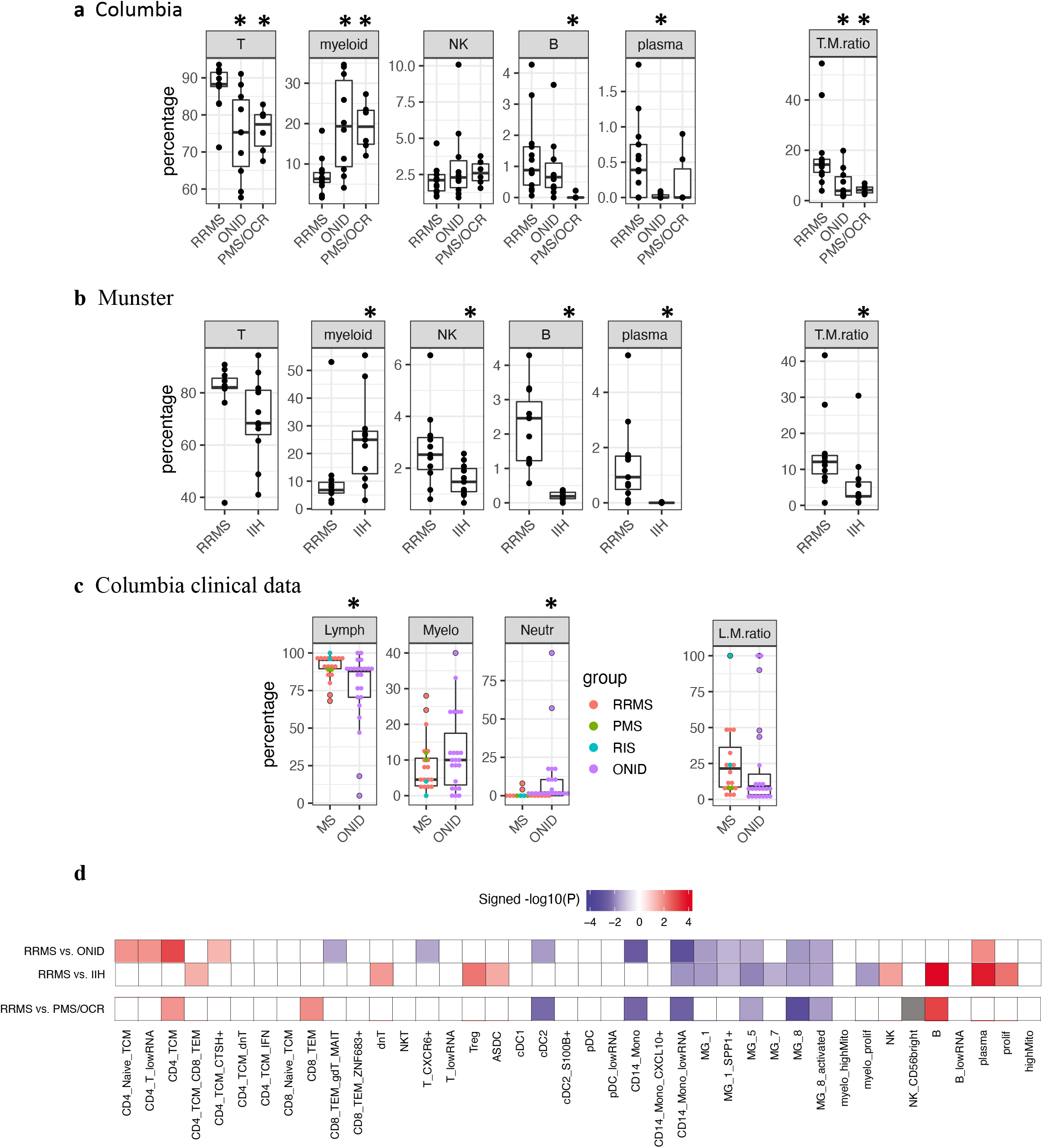

In secondary analyses, we assessed which T and myeloid subsets may be driving the observed differences in these two major cell populations in MS (*p*<0.05). As seen in **Figure 5d**, the associations with T cell subsets are modest: while CD8^+^T/gdT/MAIT and CXRC6^+^ TRM clusters are diminished in frequency, clusters containing CD4^+^ naïve, TCM, and Th17-like CD4^+^CTSH^+^ cells are increased in frequency in RRMS compared to ONID. However, this is not observed in comparison with IIH. Hence, elucidation of the role of these T cell subtypes will require further data generation from additional RRMS and ONID cases. On the other hand, the reduction in the frequencies of multiple myeloid cell subsets is more consistent between the two sets of comparisons, with several microglia-like clusters and a CD14^+^ monocyte cluster characterized by low RNA levels diminished in RRMS in comparison to both ONID and IIH. Thus, despite the limitations of having small sample sizes, we see robust results that focuses our attention on the reduction of microglia-like cells and increase in plasma cells as likely characteristic features of early RRMS.

### CSF gene expression in RRMS compared to other neuroinflammatory disorders

Having evaluated changes in the frequency of different cell populations to identify cell types that may be relevant to MS immunopathology, we undertook the complementary evaluation of gene expression in each cell subtype. Similar to our approach in differential cell frequency analysis, we performed separate comparisons between RRMS and ONID (Columbia samples) and RRMS and IIH (Munster samples) (details can be found in the **Supplementary Methods**). Genes with Bonferroni-adjusted *p*<0.05 and log2 fold change>0.25 were considered significant in each comparison in each cell type. **Supplementary Table** ◻ contain the detailed results. Several genes are found to be differentially expressed in more than 1 cell type. However, most are significant only in 1 major cell class (i.e., B, T, or myeloid cells).

105 genes were commonly differentially expressed in comparisons between RRMS and ONID, and RRMS and IIH (**Supplementary Table** ◻), and were most significantly enriched for TNF, and alpha and gamma interferon response pathways. Our findings are in line with previous literature on the enrichment of these pathways in comparisons between MS and control conditions[ref], and furthermore suggest that these identified genes and pathways are more specifically associated with MS-related processes.

Since the plasma cell population is essentially absent in ONID and IIH samples, we could not perform differential gene expression analysis for this cell type. However, we organized the results relative to this important cell population by using the Connectome^29^ pipeline to assess whether plasma cells express complementary ligands or receptors to differentially expressed genes from other CSF cell types. This could generate hypotheses on whether/how plasma cells affect or get affected by other immune cell types through ligand-receptor interactions. Our analysis found a number of upregulated ligands in T and myeloid cells in RRMS for which receptors were expressed on plasma cells (**Figure 6a**). Among these ligand-receptor pairs *CCL5-SDC1* is the most specific to plasma cells, as *SDC1* (which encodes CD138) is almost exclusively expressed in this cell type. CCL5, also known as RANTES (Regulated on Activation, Normal T cell Expressed and Secreted), is upregulated in our RRMS samples in comparison to both ONID and IIH in several CD4^+^ T cell clusters. Soluble CCL5 has been previously shown to be increased in the CSF of MS patients^30,31^ and is produced by T cells (higher expression in CD8^+^ T cells), perivascular inflammatory cells and astrocytes in MS brain tissue. It is also believed to be important in the recruitment of T cells and macrophages to MS lesions through its receptor CCR5^32^. Here, our data suggests that this chemokine might as well play an important role in the recruitment of plasma cells to the CSF in MS through its interaction with CD138.

**Figure 6.**
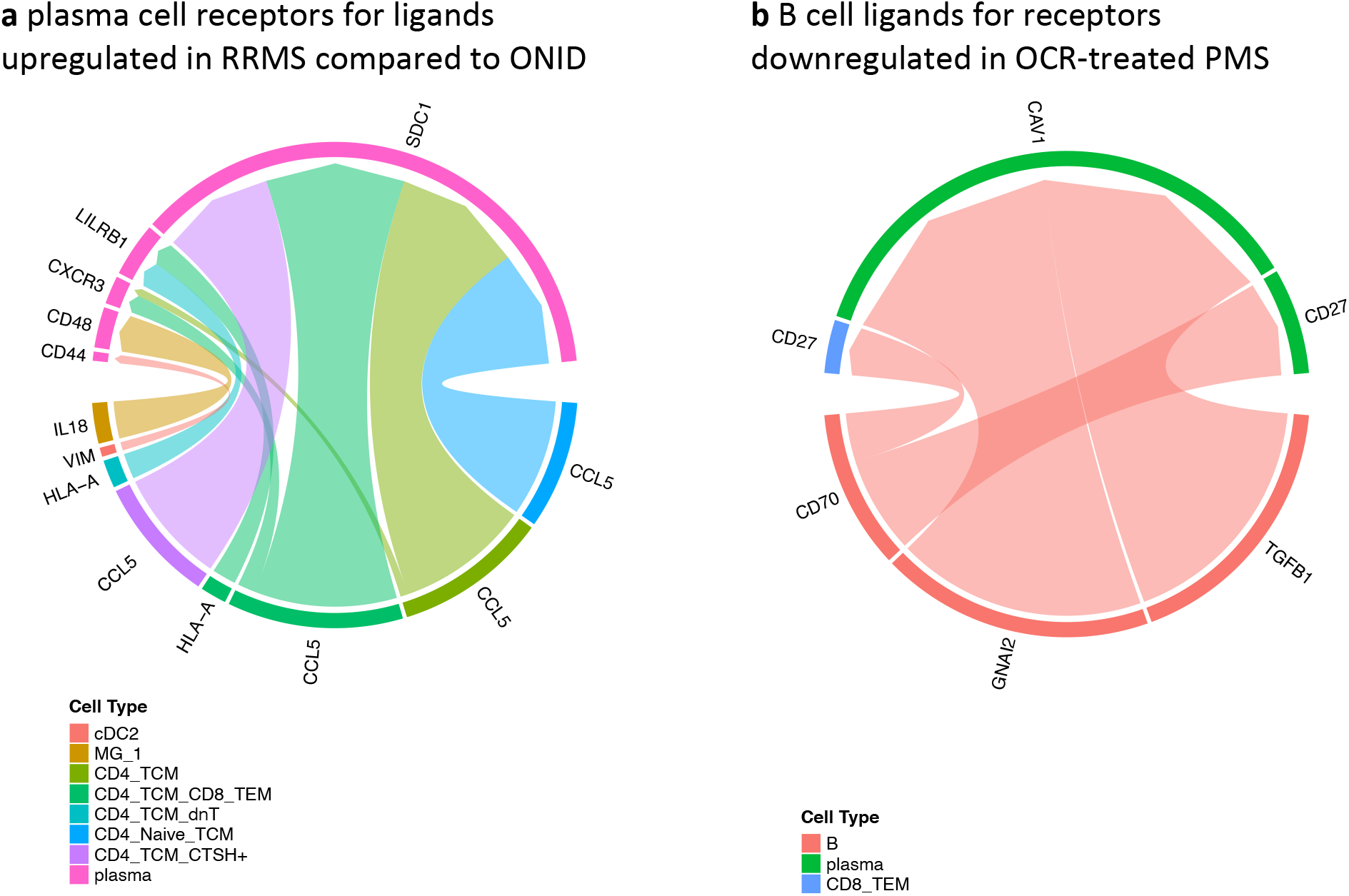

### CSF cell composition in ocrelizumab-treated MS

We also performed comparisons between RRMS and ocrelizumab (OCR)-treated progressive MS (PMS) in order to identify the likely effects of anti-CD20 therapy on CSF cells. Peripheral blood studies on the effects of ocrelizumab have demonstrated an almost complete depletion of B cells and a reduction in T cell frequency^33,34^ that remain significant during the 6-month interval between the infusions^33^. Our analyses on CSF samples from untreated RRMS and OCR-treated PMS 5 months after their last infusion recapitulate the findings from peripheral blood (**Figure 5a**): we observe significantly lower proportions of B cells in the OCR group (with 5 out of 6 OCR-treated patients having 0 B cells in CSF), in addition to significantly lower proportions of T cells and higher proportions of myeloid cells. Thus, the RRMS-related changes in these major cell classes appear to be reversed in the OCR-treated PMS participants. However, we did not observe a significant difference in the proportions of plasma cells between RRMS and our small group of OCR-treated patients. Further analyses on T and myeloid subtypes suggested that OCR-treated patients have reduction in both CD4^+^ and CD8^+^ memory T cells, while the increase in myeloid cells are seen in CD1c^+^ dendritic cells, CD14^+^ monocytes, and MG-5 and MG-8-like cells, but not in homeostatic (MG-1)-like cells (**Figure 5d**).

CD20 is expressed on almost all B cells, in addition to a smaller proportion of plasma and T (especially CD8^+^) cells^34^. Although CD20^+^ T cells are shown to be almost completely depleted in the peripheral blood of OCR-treated MS patients, it seems that the observed reduction in peripheral T cells is not fully explained by the depletion of CD20^+^ T cells^33,34^, and it is likely that more complex mechanisms are involved. Our CSF data collected 5 months after the last ocrelizumab infusion does not show a reduction in the percentage of CD20^+^ T cells in CD8-dominant T cell clusters. This might need to be interpreted in the context of the compartmentalized CSF immune environment. On the other hand, we observed a reduction in the percentage of CD20^+^ cells in the differentially abundant CD4^+^ TCM cluster (*p*=0.04) (**Supplementary Figure** ◻). Nonetheless, our analysis suggests that similar to peripheral blood, the reduction in the frequency of CSF T cells cannot be solely explained by the effect of ocrelizumab on CD20^+^ T cells.

Since our OCR-treated patients were older and had progressive MS, we cannot necessarily attribute all observed differences to ocrelizumab. However, given the profound effects of the treatment and the similarities between our major findings and blood studies, the treatment is likely to be the factor with the strongest effect, relative to age and non-relapsing disease. This is in part supported by our exploratory analysis showing no significant correlation between age and the frequency of the differentially abundant clusters among RRMS patients. Further work can leverage this observation to explore in greater detail.

### CSF gene expression in ocrelizumab-treated MS

Comparing gene expression between RRMS and OCR-treated PMS samples from the Columbia dataset, we found several genes that were differentially expressed in T, myeloid, and plasma cells (**Supplementary Table** ◻). Although a large number of genes were differentially expressed in plasma cells, we believe that the results from this cell type need to be interpreted with caution, as only two samples from the OCR-treated group contained plasma cells. Nevertheless, we have reported these results to serve as a reference for future studies. Pathways enriched in the differentially expressed genes in OCR-treated samples included the previously discussed pathways in comparisons between MS and ONID/IIH, supporting the hypothesis that anti-CD20 treatment reverses MS-related inflammatory processes.

As the B cell population was depleted in OCR-treated samples, differential gene expression analysis could not be performed for this cell type. However, we leveraged our Connectome^29^ analysis ligand-receptor association results to prioritize signaling connections between B and other CSF cells that might have been affected by B cell depletion for further investigation. Focusing on the more B cell-specific results, our findings suggested that *CD27* downregulation observed in plasma cells and CD8^+^ memory T cells in OCR-treated patients might be related to the depletion of *CD70* produced by B cells (**Figure 6b**). *CD70* encodes the sole ligand for CD27, and was expressed in 16% of B cells and smaller percentages of other cell types in our CSF samples. *CD27*, a member of the TNF-receptor superfamily, is expressed in plasma, T, B, and NK cells in our samples, and is upregulated in T cells in RRMS compared to IIH, but not ONID. Previous studies have shown that soluble CD27 level is increased in CSF of MS patients^35^, and is predictive of MS diagnosis and high relapse rate in clinically isolated syndrome (CIS)^36^. Given our findings, CD27 downregulation might play an import role in mediating the disease-modifying effects of ocrelizumab.

## Discussion

Using the largest number of CSF sc-RNAseq samples available from MS and other neuroinflammatory disorders to date, our two-stage study attempted to: (1) create an updated atlas of CSF immune cells in neuroinflammatory conditions and (2) deploy this detailed atlas to identify disease- and treatment-specific effects in MS, and prioritize CSF immune cell types, genes, pathways, and ligand-receptor interactions for further mechanistic investigations of MS and its treatment.

As a result of the larger sample size, we were able to identify novel lymphoid and myeloid cell clusters not previously described in CSF, such as CXCR6^+^ TRM and S100B^+^ cDC2, and several subclasses of CSF microglia-like cells. Given that myeloid cells on average constitute <15% of the CSF cells, we were especially more powered relative to previous studies in the detection of low frequency myeloid cell types, such as AXL^+^SIGLEC6^+^ and S100B^+^ dendritic cells, and CD14^+^CXCL10^+^ monocytes. Of note, we tried to optimize our cluster annotations by mapping our data to a comprehensive set of reference datasets. This helped increase our confidence in a number of annotations where marker genes are shared between different cell types, such as between monocytes and granulocytes. The same approach helped us better characterize the CSF microglia-like clusters by mapping CSF cell signatures to scRNA-seq data from brain microglia. Our analysis suggested that CSF cells do not represent brain microglia clusters proportionally, with the major CSF microglia-like clusters resembling brain MG-1 and MG-8, and other smaller CSF clusters resembling MG-5 and MG-7. Finally, we were able to address the question on the ontogeny of CSF microglia-like cells using data from a female participant who had received bone marrow transplant from a male donor. Our data provided causal evidence for the differentiation of CSF microglia-like cells from peripheral circulation bone marrow-derived myeloid cells, rather than from yolk-sac derived brain microglia.

Our study included samples from a variety of diagnostic groups, each of which could have large effects on the frequency of CSF cell types. Hence, we were able to cover a wide range of CSF cell composition patterns in MS and other neuroinflammatory conditions. As a result of this heterogeneity, we were more likely to find disease-, treatment-, or sample-specific effects, as were the case in our observations of CXCL13^+^ T cells specific to the lymphoma participant, plasma cell population specific to MS patients, and near complete depletion of B cells in the OCR-treated group. Using this approach, we prioritized the increase in plasma cells and the reduction in microglia-like cells as more MS-specific findings, as these changes were observed in both MS vs. ONID and MS vs. IIH comparisons. Both of these findings can be interpreted in relation to MS histopathology: it is possible that the microglia-like cells are reduced in MS CSF as they are recruited to the parenchymal lesions; and given that only a subset (71%) of MS patients had higher levels of CSF plasma cells (>0.25%), the presence of plasma cells in the CSF might be associated with MS immunopattern II^37,38^, where immunoglobulin and complement deposition are observed in active lesions. Pattern II is estimated to be the most common immunopathologic pattern in MS (56%)^37^, and patients with pattern II lesions are more likely to benefit from plasma exchange^39,40^. Given the observed heterogeneity in histopathology and response to treatment, future studies assessing whether CSF plasma cells can serve as a surrogate marker for pattern II MS lesions might help guide precision treatment in MS patients.

In addition to prioritizing CSF cell types associated with MS diagnosis (plasma and microglia-like cells, discussed above), we utilized our atlas to prioritize cell types associated with genetic susceptibility to MS. Results included a wide range of cell types, including cells involved in MS histopathology (B, plasma, microglia-like, and CD8^+^ T cells), in addition to a number of other T and myeloid cell types. Of note, contrary to the former analysis comparing MS with other neuroinflammatory conditions, the comparator group for identifying MS susceptibility genes have been healthy people. Hence, results from the MS susceptibility enrichment analysis are not necessarily specific to MS, and might include cell types susceptible to general neuroinflammatory processes shared between MS and other neuroinflammatory diseases. Nonetheless, the prioritized cell types likely have a causal role in neuroinflammation, and can be used as targets for further exploration of their specificity in divergent neuroinflammatory pathologies.

Besides genetic studies, studies of therapeutic interventions that result in the complete depletion of specific cell types can be utilized to provide causal evidence with regard to cellular dynamics and help unravel cellular interactions in *in vivo* systems. Using this concept, our study of the effects of ocrelizumab, which depletes CD20^+^ cells including all B and a small proportion of T cells from peripheral blood, provided evidence for the causal effect of this depletion on changes in the frequencies of specific lymphoid and myeloid cell subsets in MS CSF. Although the exact cellular and molecular mechanisms underlying these effects remain unknown, our ligand-receptor expression analysis prioritized CD27, expressed on CD8^+^ T and plasma cells, which downregulation might have been the direct consequence of the depletion of its sole ligand CD70 produced by B cells. Further studies are needed to assess whether these prioritized interactions contribute to the therapeutic effect of ocrelizumab. Similar approach in ligand-receptor expression analysis using data from untreated MS and ONID patients, where plasma cells are absent from non-MS CSF, prioritized CCL5 for further mechanistic analyses on whether its increased expression in MS is related to the increased frequency of CSF plasma cells, possibly through affecting its plasma cell receptor SDC1.

Our study has been able to generate new knowledge on the immune cell types present in CSF in neuroinflammatory conditions, including the introduction of new cell types, and improved characterization of previously known clusters. However, we acknowledge the limitations of our data and our analyses. Compared to CITE-seq data, where surface protein expression data is available for cells in addition to scRNA-seq data, our classification and characterization of the T cell clusters remains suboptimal. This is related to the fact that some classic functional subsets of T cells remain less distinguishable using scRNA-seq data alone^8^. We refined our cluster annotations by mapping our data to reference datasets. Although in most cases we reached a consensus by using information on cluster marker genes and annotations from multiple references, these information remain restricted by the limited cell types/states that the reference atlases include. However, using more comprehensive references have been helpful in improving the accuracy of our annotations. Although our samples were from a wide range of diagnoses, it is possible that future studies describe new disease- or treatment-specific cell types by including new diagnostic groups. Lastly, although our sample size is the largest available to date, our within-center comparisons between different diagnostic groups have relatively small sample sizes, and would benefit from sample additions in future studies.

Overall, our study provides an updated reference for future studies of CSF immune cells in neuroinflammation, along with a framework for prioritization of cell types, genes, and pathways for studies of disease conditions. Additionally, our observations on CSF cell types and genes altered in MS compared to other inflammatory disorders, and our hypotheses on molecular interactions underlying these differences can guide future studies of MS pathophysiology. Finally, our investigation of the effects of anti-CD20 therapy in CSF can serve as a proof of principle study for the utilization of scRNA-seq in assessing the effects of therapeutic interventions on cell composition and gene expression in target tissues and in generating hypotheses for the cellular and molecular mechanisms underlying those therapeutic effects.

